# Pleiotropic Mapping and Annotation Selection in Genome-wide Association Studies with Penalized Gaussian Mixture Models

**DOI:** 10.1101/256461

**Authors:** Ping Zeng, Xinjie Hao, Xiang Zhou

## Abstract

**Motivation:** Genome-wide association studies (GWASs) have identified many genetic loci associated with complex traits. A substantial fraction of these identified loci are associated with multiple traits – a phenomena known as pleiotropy. Identification of pleiotropic associations can help characterize the genetic relationship among complex traits and can facilitate our understanding of disease etiology. Effective pleiotropic association mapping requires the development of statistical methods that can jointly model multiple traits with genome-wide SNPs together.

**Results:** We develop a joint modeling method, which we refer to as the integrative MApping of Pleiotropic association (iMAP). iMAP models summary statistics from GWASs, uses a multivariate Gaussian distribution to account for phenotypic correlation, simultaneously infers genome-wide SNP association pattern using mixture modeling, and has the potential to reveal causal relationship between traits. Importantly, iMAP integrates a large number of SNP functional annotations to substantially improve association mapping power, and, with a sparsity-inducing penalty, is capable of selecting informative annotations from a large, potentially noninformative set. To enable scalable inference of iMAP to association studies with hundreds of thousands of individuals and millions of SNPs, we develop an efficient expectation maximization algorithm based on an approximate penalized regression algorithm. With simulations and comparisons to existing methods, we illustrate the benefits of iMAP both in terms of high association mapping power and in terms of accurate estimation of genome-wide SNP association patterns. Finally, we apply iMAP to perform a joint analysis of 48 traits from 31 GWAS consortia together with 40 tissue-specific SNP annotations generated from the Roadmap Project. iMAP is freely available at www.xzlab.org/software.html.

## 1. Introduction

Genome-wide association studies (GWASs) have identified thousands of genetic variants associated with many complex diseases and traits (MacArthur, et al., 2017). A substantial fraction of these identified loci often display association with more than one trait — a phenomenon known as pleiotropy (Solovieff, et al., 2013). Notable examples of pleiotropic genes include *PTPN22* and *HLA-DRB1* that are associated with several different autoimmune disorders (The Wellcome Trust Case Control Consortium, 2007); *CLPTM1L* with multiple types of cancers (Fletcher and Houlston, 2010); *CACNA1C* with bipolar disorder and schizophrenia (Cross-Disorder Group of the Psychiatric Genomics Consortium, 2013); *ANGPTL3*, *TIMD4*, *PPP1R3B* and *TRIB1* with multiple plasma lipid traits (Teslovich, et al., 2010); *MYL2* with various metabolic traits (Kim, et al., 2011); *ZFAT* with total cholesterol, blood pressure and cardiovascular disease (Smith, et al., 2010; Warren, et al., 2017); and ABO with blood group, various blood measurements and coronary artery disease (Bulik-Sullivan, et al., 2015; Pickrell, et al., 2016). Overall, it has been estimated that 4.6% of the previously identified associated variants and 16.9% of the previously identified associated genes show pleiotropic effects (Sivakumaran, et al., 2011). The percentage of pleiotropic association is often much higher in biologically relevant traits. For example, about 44%∼70% of genetic variants that are associated with one autoimmune disease are also associated with another (Cotsapas, et al., 2011; Jostins, et al., 2012; Solovieff, et al., 2013). In addition, joint heritability analyses of multiple traits have also revealed substantial genetic correlation among various psychiatric disorders (Bulik-Sullivan, et al., 2015; Cross-Disorder Group of the Psychiatric Genomics Consortium, 2013), between schizophrenia and brain anatomic measurements (Lee, et al., 2016) or amyotrophic lateral sclerosis (McLaughlin, et al., 2017), between neuropsychiatric disorders and several metabolic traits (Lane, et al., 2017), as well as among multiple autoimmune diseases (Ji, et al., 2017; Zhernakova, et al., 2009).

Because of the prevalence and importance of pleiotropy, many statistical methods have been developed to identify genetic variants that are associated with multiple phenotypes (Li and Kellis, 2016; Liu, et al., 2016; Solovieff, et al., 2013; van der Sluis, et al., 2013; Zhou and Stephens, 2014). Most of these methods rely on multivariate models to analyze multiple traits jointly. Unlike the commonly used univariate methods that examine one trait at a time, multivariate methods analyze multiple traits together while properly accounting for the phenotypic correlation among them. As a result, multivariate methods often substantially improve association mapping power compared with univariate methods. For example, compared with univariate analysis, multivariate analysis achieves ∼50% power gain in mapping systolic blood pressure and other comorbid traits (Andreassen, et al., 2014) as well as in mapping multiple correlated blood lipid traits (Stephens, 2013; Zhou and Stephens, 2014). Applications of multivariate methods in GWASs have identified a large number of pleiotropic associations (He, et al., 2013; Solovieff, et al., 2013; Van der Sluis, et al., 2015; Zhu, et al., 2015).

Among the existing methods for pleiotropic association mapping, mixture models have attracted substantial recent attention. Mixture models attempt to classify genome-wide single nucleic polymorphisms (SNPs) into different categories based on their effect sizes on the jointly analyzed traits. For example, the Bayesian co-localization test (Giambartolomei, et al., 2014) and its recent extension and implementation in the computational tool, gwas-pw (Pickrell, et al., 2016) (pair-wise traits analysis of GWAS), analyzes pairs of traits jointly. In its simplified version, gwas-pw divides SNPs into those that are associated with both traits (i.e. pleiotropic associations), those that are associated with only one trait, and those that are associated with neither. By partitioning SNPs into different association categories and inferring the genome-wide association pattern, mixture models can improve association mapping power (Andreassen, et al., 2014; Giambartolomei, et al., 2014). In addition, by computing the proportion of SNPs in each category, mixture models have the potential to provide evidence supporting potential directional causality between the two analyzed phenotypes (Pickrell, et al., 2016). Specifically, if SNPs associated with the first trait are also associated with the second trait, but not vice versa, then the first trait may causally influence the second trait. Therefore, mixture model methods can facilitate the identification and interpretation of pleiotropic associations and can help better characterize the genetic relationship among traits (Chen, et al., 2016; Ernst and Kellis, 2010; Ernst, et al., 2011; Fu, et al., 2014; Ionita-Laza, et al., 2016; Lonsdale, et al., 2013; Lu, et al., 2016).

Mixture models are recently extended to integrate external SNP functional annotations to further improve power of pleiotropic mapping. For example, GPA (Genetic analysis incorporating Pleiotropy and Annotation) (Chung, et al., 2014) and its gene-set analysis extension EPS (Empirical Bayes approach for integrating Pleiotropy and tissue-Specific information) (Liu, et al., 2016), extend mixture models by jointly modeling SNP association pattern together with the distribution of the SNP functional annotations. SNP functional annotations are developed to characterize the function of genetic variants (Dixon, et al., 2015; Kellis, et al., 2014; Lonsdale, et al., 2013). For example, we can now classify genetic variants based on their genomic location (e.g. coding, intron and intergenic variants), role in protein structure and function (e.g. SIFT score (Kumar, et al., 2009) or PolyPhen score (Adzhubei, et al., 2013)), ability to regulate gene expression (e.g. eQTL and ASE evidence (Pickrell, et al., 2010; Tung, et al., 2015)), biochemical function (e.g. DNase I hypersensitive sites, metabolomic QTL evidence and chromatin states (Ernst and Kellis, 2012; McVicker, et al., 2013; Pique-Regi, et al., 2011)), evolutionary significance (e.g. GERP score (Cooper, et al., 2005)), and/or a combination of all these annotations (e.g. CADD score (Kircher, et al., 2014) and Eigen score (Ionita-Laza, et al., 2016)). These functional annotations can be important predictors for SNP effects. Studies have shown that SNPs in certain functional categories (e.g. in promoters and enhancers) are more likely to be causal (Pickrell, 2014; Schork, et al., 2013), tend to have larger effect sizes, and explain more heritability than SNPs in other categories (e.g. introns) (Gusev, et al., 2014; Kichaev, et al., 2014). Therefore, by integrating functional annotations into pleiotropic association mapping, GPA can identify enriched functional annotations for each SNP association category, with which to further improve association mapping power (Chung, et al., 2014).

Despite the promising results from mixture models for pleiotropic association mapping, existing mixture methods have two important limitations. First, perhaps rather surprisingly, existing mixture methods do not account for phenotypic correlation among jointly analyzed traits and effectively assume phenotypic independence. Given that explicit modeling of phenotypic correlation is the key for effective association mapping and phenotype/risk prediction of pleiotropic traits (Hu, et al., 2017; Maier, et al., 2015; Stephens, 2013), assuming phenotype independence likely reduces the effectiveness of existing mixture models. Second, existing mixture models are only able to integrate binary annotations and can only handle a relatively small number of annotations. Analyzing only a few binary annotations may fail to take advantage of the increasingly large number of annotations that are being generated today, among which many are continuous (Ionita-Laza, et al., 2016; Kircher, et al., 2014; Li and Kellis, 2016; Roadmap Epigenomics Consortium, et al., 2015; The ENCODE Project Consortium, 2012).

Here, we develop a novel mixture modeling method, which we refer to as the **i**ntegrative **MA**pping of **P**leiotropic association (iMAP), for association mapping of pair-wise traits. iMAP relies on a multinomial logistic regression model to incorporate a large number of binary and continuous SNP annotations, and, with a sparsity-inducing penalty term, is capable of selecting a small, informative set of annotations. In addition, iMAP directly models summary statistics from GWASs and uses a multivariate Gaussian distribution to account for phenotypic correlation between traits. As a result, iMAP is capable of integrating both binary and continuous SNP annotations, selecting informative annotations from a large set of potentially noninformative ones, and using GWAS summary statistics while simultaneously accounting for phenotypic correlation between traits. We also adopt a recently developed efficient approximation algorithm for penalized regression (Wang and Leng, 2007; Zeng, et al., 2014) to enable scalable inference in iMAP. With simulations, we illustrate the benefits of iMAP both in terms of improving association mapping power and in terms of accurate estimation of SNP association patterns. Finally, we apply iMAP to analyze 48 traits from 31 GWAS consortium studies together with 40 tissue-specific SNP annotations from the Roadmap epigenomics project (Roadmap Epigenomics Consortium, et al., 2015). iMAP is freely available at www.xzlab.org/software.html.

## 2. Methods

We provide an overview of iMAP here, details are available in Text S1. Our main presentation of iMAP will focus on analyzing pairs of traits that are measured on the same set of individuals from a GWAS. Extensions of our method to situations where the two traits come from different GWASs are trivial and are presented in Text S1. Extensions of our method to analyzing more than two traits are not straightforward, with potential statistical and computational complications, and will be discussed later in the Discussion. For each SNP in turn, we consider the following bivariate model

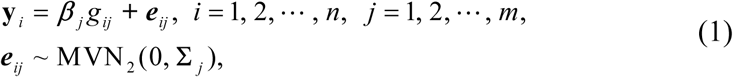

where *n* is the number of individuals; *m* is the number of SNPs; **y**_i_ is a two-vector of phenotypes for individual *i*; *g*_*ij*_ is the genotype for SNP *j* of individual *i*; ***β***_*j*_ is the corresponding two-vector of effect sizes of SNP *j* on the two phenotypes; ***e***_*ij*_ is a two-vector of residual errors that follow a bivariate normal distribution with a two by two covariance matrix **Σ**_*j*_; and MVN_2_ denotes a two-dimensional multivariate normal distribution (i.e. bivariate normal distribution). iMAP naturally accounts for phenotypic correlation by modelling the covariance matrix **Σ**_*j*_. While Equation (1) is primarily aimed to deal with quantitative traits, we also use Equation (1) to model binary traits by treating binary data as continuous values following many previous studies (Moser, et al., 2015; Speed and Balding, 2014; Weissbrod, et al., 2016; Zhou, et al., 2013). Methodologically, modeling binary data with linear model can be justified by the fact that a linear model is a first order Taylor approximation to a generalized linear model; and the approximation is accurate when the SNP effect size is small (Zhou, et al., 2013) - a condition that generally holds as most complex phenotypes are polygenic and are affected by many SNPs with small effects (Visscher, et al., 2017).

The above model is specified on each SNP *j*. To infer genome-wide pleiotropic association pattern and improve association mapping power, following (Chung, et al., 2014; Pickrell, et al., 2016), we model all SNPs jointly by assuming that the joint likelihood for all SNPs is simply a product of the likelihood for each SNP. The joint likelihood approximation by the product of individual likelihoods can be justified from a composite likelihood perspective (Larribe and Fearnhead, 2011; Varin, et al., 2011) (Text S1). To facilitate information sharing across genome-wide SNPs, we assign a common prior on the effect sizes and assume that each ***β***_*j*_ *a priori* follows a four-component Gaussian mixture distribution

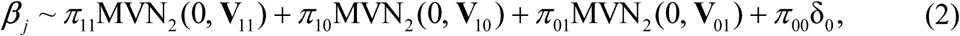

with prior probabilities *π*_*k*_ (*k* = 11, 10, 01 and 00) that sum to one. Here *π*_11_ represents the prior probability that a SNP is associated with both traits; 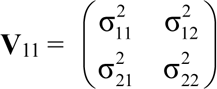 two by two covariance matrix modeling the covariance of SNP effects on the two traits. *π*_10_ represents the prior probability that a SNP is associated with the first trait but not the second; 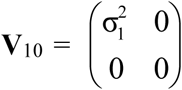 is a low-rank covariance matrix restricting that only the effect size for the first trait is nonzero. *π*_01_ represents the prior probability that a SNP is associated with the second trait but not the first; 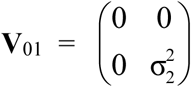 is a low-rank covariance matrix restricting nonzero effect only for the second trait. Finally *π*_00_ represents the prior probability that a SNP is not associated with any traits; *δ*_0_ denotes a point mass at zero. We specify conjugate priors for the three hyper-parameters V and we borrow information across genome-wide SNPs to infer these parameters (Text S1). The estimated probability of a SNP having nonzero effects on any traits (i.e. *π*_11_ + *π*_10_ + *π*_01_) is commonly referred to as the posterior inclusion probability (PIP), which represents association evidence for the given SNP. We use PIP to prioritize SNPs in the present study. We also use PIP to provide a conservative estimate of false discovery rate (FDR) (Benjamini and Hochberg, 1995) based on local false discovery rate (Efron, et al., 2001) via the direct posterior probability approach (Newton, et al., 2004). FDR is a common approach applied for multiple testing settings. Effective control of FDR ensures that the proportion of false discoveries among the identified SNP associations is kept in check regardless of the number of tests performed. However, we also note that an FDR of 0.05 can be less stringent than the usual genome-wide p-value threshold of 5×10^−8^ that aims to control for a family-wise error rate (FWER) of approximately 0.05.

To increase association mapping power and facilitate association results interpretation, we incorporate functional SNP annotations from external data into the above model. To do so, we assume that all SNPs are characterized by the same set of *q* annotations. For SNP *j*, we denote **x**_*j*_ = (1, *x*_*j*1_, *x*_*j*2_,…, *x*_*jq*_)^T^ as a (*q +* 1)-vector of annotation values that include a value of one to represent the intercept. These annotations can be either discrete or continuous. For example, one annotation could be a binary indicator on whether a SNP resides in exonic regions, while another annotation could be a continuous value of the CADD (Kircher, et al., 2014) or Eigen (Ionita-Laza, et al., 2016) score for the given SNP. To simplify presentation, we assemble the annotation vectors across all SNPs into an *m* by (*q* + 1) annotation matrix **X**, where each row of **X** contains the annotation vector for the corresponding SNP. We then link the annotation matrix **X** to the mixture probabilities (*π*_11_, *π*_10_, *π*_01_ and *π*_00_) through a multinomial logit (mlogit) regression model

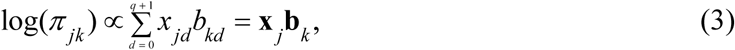

where *k* = 11, 10, 01 or 00; and each **b**_*k*_ = (*b*_0_, *b*_1_, *b*_2_,…, *b*_q_) is a (*q* + 1)-vector of annotation coefficients that include an intercept. We choose *k* = 00 as the reference category and set **b**_00_ = **0** to ensure model identifiability. Note that, in the case where SNPs belong to only two categories (e.g. causal vs non-causal), the mlogit model reduces to a logistic regression model that has been commonly used to link functional annotations to SNP causality (Carbonetto and Stephens, 2013; Wen, et al., 2016; Wen, et al., 2015). We use the mlogit model here because it naturally extends the logistic model to cases where SNPs can belong to more than two categories (e.g. four categories here: *k* = 11, 10, 01 and 00).

With the growing number of SNP annotations nowadays, however, it becomes increasingly challenging to model all annotations in the above mlogit model. Examining one annotation at a time (Kichaev, et al., 2014; Pickrell, 2014) does not take full advantage of the correlation structure among annotations and may fail to properly account for multiple testing issue (Chen, et al., 2016). While including all annotations jointly without any prior assumption may incur heavy computational burden, reduce the degrees of freedom, and lead to a potential loss of power. Here, to handle a large number of annotations, we hypothesize that only a fraction of these annotations are likely informative. Subsequently, we impose a penalty term 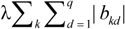 onto the log likelihood of model (3) to select the important annotations via Lasso (Tibshirani, 1996) (Text S1; *λ* is the tuning parameter).

We develop an Expectation-Maximization (EM) algorithm for parameter inference in iMAP. Briefly, we view the mixture group assignment of each SNP as missing data and impute them in the expectation step. In the maximization step, we adopt the recently developed computational strategy for the mlogit model (Hasan, et al., 2016) to take full advantage of the sparse structure of the Hessian matrix, in order to estimate the annotation parameters **b** in a computationally efficient fashion. In addition, we employ an efficient approximation algorithm for lasso based on the least square approximation (LSA) (Wang and Leng, 2007; Zeng, et al., 2014) to further improve computational efficiency. We rely on the Bayesian information criterion (BIC) to select informative annotations. Details of our inference algorithm are provided in Text S1.

Finally, we note that, while our main presentation of the model and algorithm rely on individual-level phenotypes and genotypes, iMAP can be fitted using only summary statistics in terms of marginal **z** scores for each phenotype (Text S1). For example, the phenotypic covariance matrix **Σ** can be estimated using the sample covariance of marginal z scores from null SNPs (Stephens, 2013). The EM algorithm can also be fitted using marginal **z** scores only. Therefore, in the following simulations, while we use individual-level genotypes to simulate phenotypes, we obtain marginal **z** scores afterwards to fit iMAP. In the real data application, we also use marginal **z** scores to fit iMAP.

## 3. Results

### 3.1 Simulation results

#### 3.1.1 Power to identify associated SNPs

We first performed simulations to compare iMAP with other methods in terms of power to detect true associations. Simulations largely follow previous studies (Chung, et al., 2014) with details provided in Text S2. Briefly, we obtained 15,495 independent SNPs in 10,000 individuals from the GERA study (Banda, et al., 2015). Among these SNPs, we considered 500 to be causal for each trait, generated causal effects independently from a normal distribution, and summed simulated residual errors to form the two traits. Due to the small number of individuals, to ensure sufficient power, our main simulations considered cases where the proportion of phenotypic variance of each phenotype explained (PVE) (Zhou, et al., 2013) by all causal SNPs is 0.50. However, in some scenarios, we also performed additional simulations to examine cases where the PVE by all causal SNPs is 0.20. To allow for pleiotropic associations, these 500 causal SNPs were selected in a way that a certain proportion of them (in the range of 0 ∼ 100%, with 20% increments) have nonzero effects on both traits. To allow for phenotypic correlation, the residual errors were generated from a bivariate normal distribution with a fixed covariance. Besides phenotypes, we also simulated two sets of continuous SNP annotations: an informative set where annotation values are dependent on SNP causality and another noninformative set where annotations do not depend on SNP causality. Overall, the simulation strategy employed here substantially deviates from our own modeling assumption; instead, it largely follows the modeling assumption employed in GPA (Chung, et al., 2014). With the genotypes and simulated phenotypes, we obtained marginal z scores and paired them with SNP annotation information to run analysis. We performed 100 simulation replicates for each setting. We considered four different methods for comparisons (Text S1): (i) univariate analysis that fits a linear model on each trait separately; (ii) gwas-pw (Pickrell, et al., 2016); (iii) GPA (Chung, et al., 2014); and (iv) iMAP. We computed the power to detect causal SNPs at the false discovery rate (FDR) of 0.05 (or 0.10).

We consider three sets of simulations. We performed the first set of simulations to illustrate the benefits of modeling phenotypic correlation between traits. To do so, we varied the residual error covariance between the two simulated traits from −0.8 to 0.8 (= −0.8, −0.5, −0.2, 0, 0.2, 0.5, 0.8) to introduce negative or positive phenotypic correlations. We did not include any annotations in simulations here to exclude the influence of annotation on power comparison. In the simulations, all methods provide slightly conservative estimates of false discovery rate (FDR) at the given level of 0.05 or 0.10 (Figure S1), suggesting proper control of FDR by these methods. We also show power of different methods at a fixed FDR of 0.05 for positive phenotypic correlations in Figure 1A. The corresponding results for FDR of 0.10 are shown in Figure S2, while the corresponding results with negative phenotypic correlations are shown in Figure S3 and S4; both sets of results are similar to Figure 1A. Based on Figure 1A, we also contrast iMAP with GPA and display the power gain of iMAP in Figure 1B. In terms of power, both the proportion of pleiotropic effects and phenotypic correlation influence the relative power among methods. First, in the absence of pleiotropic effects, the power of iMAP and GPA are comparable with each other and with that of univariate analysis. However, both iMAP and GPA outperform the univariate analysis in the presence of pleiotropic effects. Second, when phenotypes are independent, the power of iMAP and GPA are comparable with each other. However, with increasing phenotypic correlation, the power of iMAP improves while the power of GPA (and gwas-pw) decays, highlighting the importance of explicit modeling of phenotypic correlation. We notice that gwas-pw is often underpowered when compared to GPA and iMAP, even if we performed simulations according to the gwas-pw study (Text S2; Figure S5). We attribute the relatively low power of gwas-pw to the fact that gwas-pw was not originally designed as an association mapping method.

**Fig.1.**
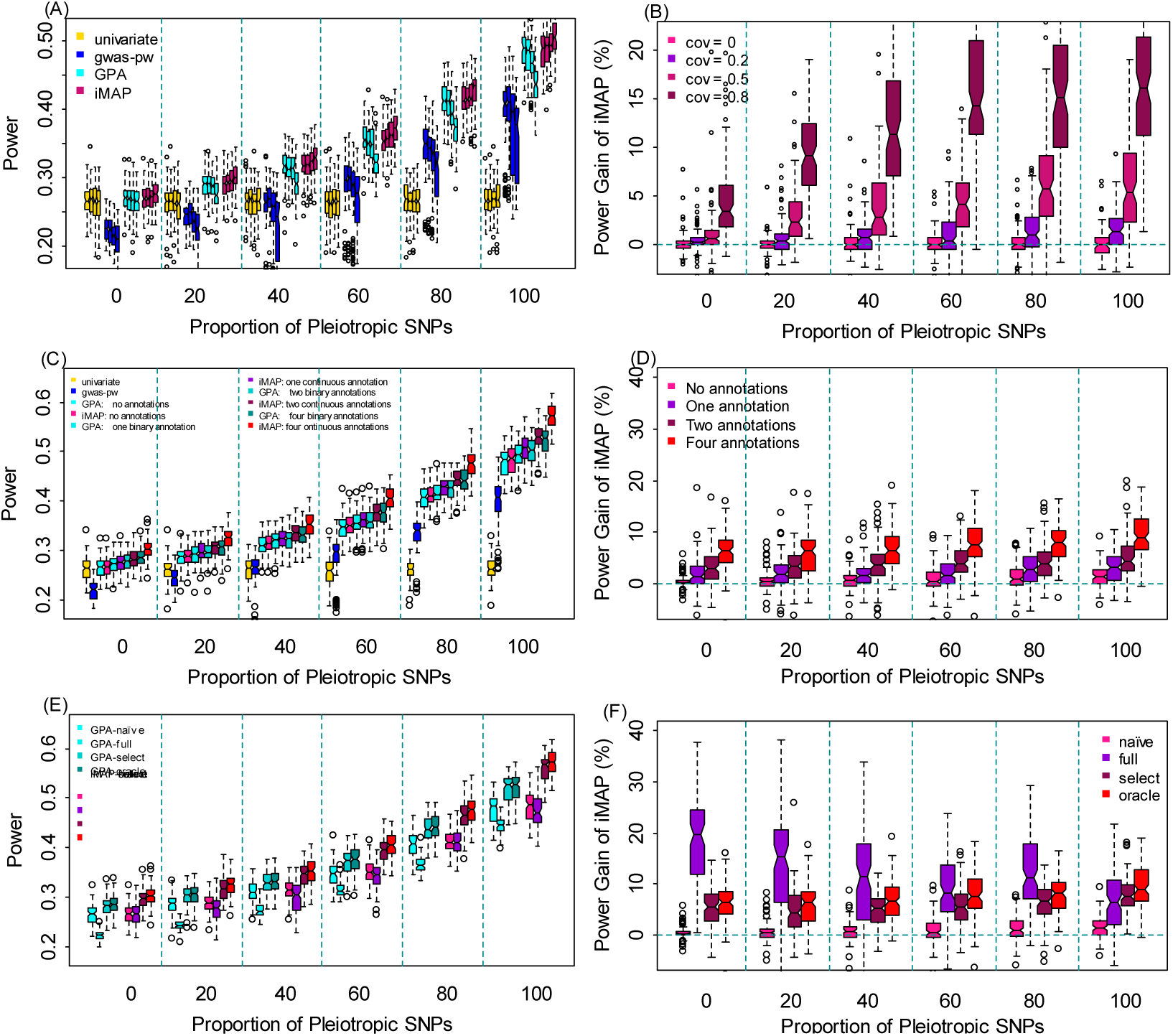
Comparison of power in detecting associated SNPs by various methods in simulations. In all simulations, the proportion of pleiotropic causal SNPs varies from 0% to 100% (x-axis). Power is measured at a fixed false discovery rate (FDR) of 0.05. (A) Power of four methods (univariate analysis, gwas-pw, GPA and iMAP) in the setting where the two traits are positively correlated. For each method, the four boxplots at each pleiotropic proportion level correspond to four different phenotypic covariance values of 0, 0.2, 0.5 and 0.8, respectively. (B) Power gain of iMAP with respect to GPA computed based on panel (A). (C) Power of four methods (univariate analysis, gwas-pw, GPA and iMAP) in the presence of informative annotations. Variations of GPA and iMAP that incorporate a different number of annotations (0, 1, 2 and 4) are also presented. (D) Power gain of iMAP with respect to GPA computed based on panel (C). (E) Power of GPA and iMAP in the presence of four informative annotations and 100 noninformative annotations. Different variations of GPA and iMAP are considered: the naïve version does not incorporate any annotations; the full version includes all the annotations; the select version performs annotation selection; and the oracle version uses the four informative annotations. (F) Power gain of iMAP with respect to GPA computed based on panel (E).

We performed the second set of simulations to illustrate the benefits of using the original continuous SNP annotations versus dichotomizing them into binary ones. To do so, we simulated four sets of informative SNP annotations that all have continuous values. We used these continuous annotations directly for iMAP but dichotomized them into binary annotations for GPA, as GPA can only take binary annotations. For both GPA and iMAP, we considered settings where 0-4 annotations were incorporated into the model. To minimize the influence of phenotypic correlation on comparison, we set the residual error covariance to be 0 and used independent traits. The power of different methods at FDR=0.05 are shown in Figure 1C. The corresponding results for FDR=0.10 are shown in Figure S6. We also contrast iMAP with GPA and display the power gain of iMAP in Figure 1D. Again, all methods provide reasonably and slightly conservative estimate of FDR (Figure S7). In terms of power, GPA and iMAP perform similarly as the univariate analysis in the absence of annotations and pleiotropic SNPs. Both GPA and iMAP outperform the univariate analysis in the presence of pleiotropic SNPs or when annotations are available. Importantly, because iMAP models the original continuous annotations directly, iMAP is slightly more powerful than GPA in the presence of annotations and its power gain increases with increasing number of informative annotations. For example, when the proportion of pleiotropic SNPs is 20%, the average power gain of iMAP versus GPA is 0.371%, 1.82%, 3.43% or 6.12%, when 1-4 annotations are incorporated, respectively (Figure 1D). The power difference between iMAP and GPA is mainly due to the use of continuous annotations, as iMAP and GPA are comparable to each other when the same set of binary annotations were used (Figure S8). Besides the main setting where the PVE by all causal SNPs was set to be 0.5, we also examined small effect settings where the PVE by all causal SNPs was set to be only 0.2. The power of all the compared methods are low in this setting, though the rank among the compared methods remains the same (Figure S9).

Finally, we performed a third set of simulations to illustrate the benefits of annotation selection when a large number of annotations are present. To do so, we used the same four informative annotations above and simulated a large number (100) of noninformative annotations whose values were independent of SNP causality. Again, we set the residual error covariance to be 0 and used independent traits. iMAP analyzes all annotations jointly through a penalized regression framework to select informative ones among them. To better understand the effectiveness of annotation selection, besides the standard iMAP, we also considered three variations of iMAP in method comparison: (i) an iMAP-naïve method that does not incorporate any annotation; (ii) an iMAP-full method that naively incorporates all annotations without selection; and (iii) an iMAP-oracle method that uses only the four informative annotations. For additional comparison, we also considered different ways of applying GPA: (i) GPA-naïve that does not include any annotation; (ii) GPA-select that examines one annotation at a time, computes for each annotation a likelihood ratio statistic, and uses a likelihood ratio statistic cutoff of 12 (which corresponds to an approximate p value of 5.0×10^−4^) to select a set of informative annotations that are further included into the GPA model for a final analysis; note that the original GPA method does not provide this option; (iii) GPA-full that includes all annotations to the model; and (iv) GPA-oracle where the correct four informative annotations are supplied. Note that the oracle version of GPA and iMAP represents an upper limit of power for the two methods. Here we did not consider the univariate analysis and gwas-pw as the two methods cannot handle annotations and were generally less powerful compared to GPA and iMAP. We display the power comparison results among different variations of iMAP and GPA in Figure 1 E-F and Figure S10. A few patterns are obvious. First, in the presence of a large number of annotations, approaches that naively model all annotations together (i.e. iMAP-full and GPA-full) are often much less powerful compared to approaches that do not consider annotations at all (i.e. iMAP-naïve and GPA-naïve), suggesting that the small degree of freedom in the full model impairs model performance. Second, approaches that select annotations (i.e. iMAP and GPA-select) almost always result in a great power gain compared to naïve approaches (i.e. iMAP-naïve and GPA-naïve), and can often achieve similar power as the corresponding oracle models (i.e. iMAP-oracle and GPA-oracle). The competent performance of selection approaches highlights the importance of performing annotation selection. Importantly, even with annotation selection, iMAP and GPA still provide effective estimate of FDR (Figure S11). Finally, iMAP relies on the formal Lasso penalty to perform annotation selection and is more powerful than GPA-select which relies on a simpler procedure to select annotations (Figure 1F). The comparative results between iMAP and GPA-select is consistent with early literature on Lasso being more effective compared with the subset selection approach (Tibshirani, 1996; Zeng, et al., 2014; Zou, 2006). Importantly, iMAP is capable of selecting the informative annotations with high precision and can accurately estimate the annotation coefficients (Table 1 and Figure S12). While our main simulation setting uses four independent annotations with relatively large effects, we also examined a setting where we include ten independent annotations with relatively small effects along with another 100 null annotations. As expected, because of small annotation effects, the accuracy to select the true annotations in this setting reduces substantially. However, the relative performance of various methods remains the same, with iMAP greatly outperforming iMAP-full (Table S2). We also assessed the performance of iMAP in a four-annotations setting where the first two annotations are correlated with each other and the second two annotations are also correlated with each other, with correlation coefficient varying from 0.2 to 0.8. The 100 null annotations are independent each other and are not correlated to the four true annotations. When correlation is low or moderate (e.g. 0.2 or 0.5), the accuracy of various methods reduces slightly with ranking among them remaining the same. In contrast, when correlation is high (e.g. 0.8), while iMAP outperforms iMAP-full in most cases, it can occasionally perform worse than iMAP-full (Table S3). The results with correlated annotations are consistent with those observed previously, and can be improved when by replacing the selection method Lasso with, for example, Elastic Net (Zou and Hastie, 2005).

**Table 1.** Accuracy of iMAP in selecting informative annotations and in estimating the annotation coefficients in the setting where four independent annotations with relatively large effects are present. Note: Simulations were carried out in the presence of four informative annotations and 100 non-informative annotations for various proportion of pleiotropic causal SNPs (rows). The “True” column lists the number of selected correct non-zero annotation parameters inside the mlogit model. Note that a total of 12 (= 3 × 4) non-zero annotation parameters are expected in the presence of four informative annotations. The “False” column lists the number of selected incorrect non-zero annotation parameters. MSE denotes the median squared error for the estimated annotation parameters across 100 simulation replicates for three different versions of iMAP: the oracle version uses the four informative annotations; the select version performs annotation selection; and the full version includes all annotations without selection.

| Proportion of pleiotropic<br>causal SNPs (%) | True | False | MSE |  |  |
| --- | --- | --- | --- | --- | --- |
|  |  |  | Oracle | Select | Full |
| 0 | 3.53 | 0.38 | 0.05 | 0.28 | 383.52 |
| 20 | 7.81 | 0.53 | 0.41 | 2.38 | 300.15 |
| 40 | 9.72 | 1.20 | 0.29 | 2.03 | 68.63 |
| 60 | 5.77 | 0.81 | 0.31 | 2.58 | 12.53 |
| 80 | 4.98 | 0.75 | 0.56 | 2.70 | 17.46 |
| 100 | 3.15 | 0.37 | 0.12 | 2.17 | 18.38 |

#### 3.1.2 Estimating causal SNP proportions and annotation coefficients

A key feature of mixture models is that they can provide estimates for the proportion of SNPs that have various effects on the two phenotypes. These proportion estimates can help shed light on the genetic architecture of complex traits. In terms of iMAP, we are often interested in estimating: *π*_11_, the proportion of pleiotropic SNPs that are associated with both traits; *π*_10_ (or *π*_01_), the proportion of SNPs that are associated with only one trait; and *π*_11_/(*π*_10_+*π*_11_) (or *π*_11_/(*π*_01_+*π*_11_)), the proportion of SNPs associated with one trait that are also associated with the other. The last quantity has been used to evaluate causality of one trait on the other (Pickrell, et al., 2016). Specifically, a large *π*_11_/(*π*_10_+*π*_11_) and a small *π*_11_/(*π*_01_+*π*_11_) suggest that a large fraction of SNPs associated with the first trait is also associated with the second trait, but not vice versa, indicating that the first trait may causally affect the second trait. A small *π*_11_/(*π*_10_+*π*_11_) and a large *π*_11_/(*π*_01_+*π*_11_) indicate that the second trait may causally affect the first trait. On the other hand, a large *π*_11_/(*π*_10_+*π*_11_) and a large *π*_11_/(*π*_01_+*π*_11_) indicate that both traits may share common biological pathways. While we caution against the causal interpretation of *π*_11_/(*π*_10_+*π*_11_) (or *π*_11_/(*π*_01_+*π*_11_)) in association studies, we do consider *π*_11_/(*π*_10_+*π*_11_) (or *π*_11_/(*π*_01_+*π*_11_)) as a useful quantity to be estimated along with the original parameters *π*_11_, *π*_10_ and *π*_01_. To compare method performance in estimating the above quantities, we focused on the second simulation setting described in the previous section and applied three different methods (gwas-pw, GPA and iMAP) to examine their performance in the presence of annotations. For better visualization, we contrast the estimated values with the true values and show their differences in Figure 2A-C. Overall, iMAP generates estimates that are slightly closer to the truth than either GPA or gwas-pw across a range of pleiotropic SNP proportions and various numbers of annotations. The slightly accuracy gain in iMAP is likely due to its higher power in identifying SNP associations.

**Fig.2.**
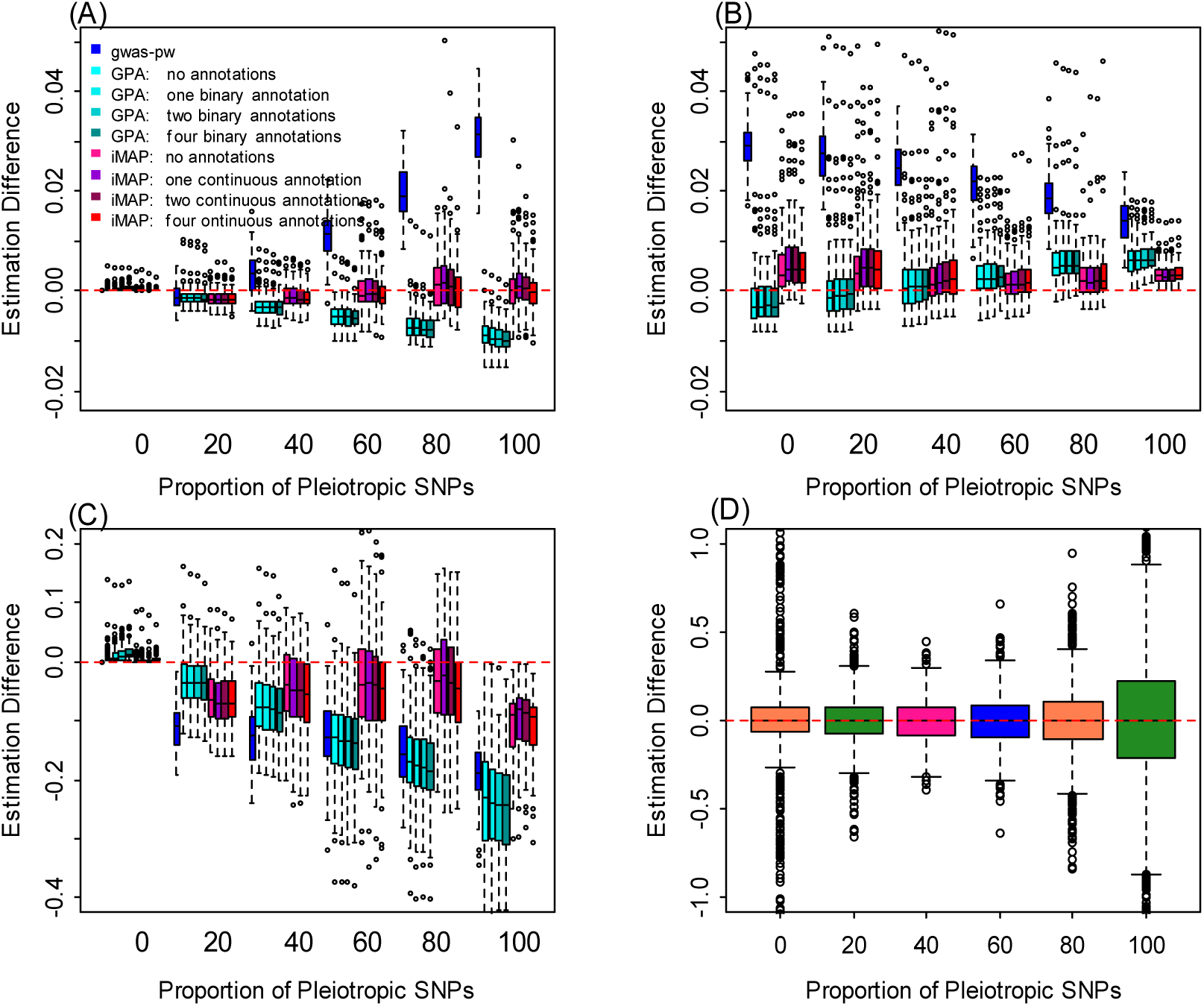
Estimation accuracy for the proportions of different SNP association categories by different methods in simulations. Methods for comparison include gwas-pw, GPA and iMAP. Because four informative annotations are present, we considered variations of GPA and iMAP that incorporate a different number of annotations (0, 1, 2 and 4). The difference between the estimated values and truth (y-axis) are computed for various settings where the proportion of pleiotropic causal SNPs varies from 0% to 100% (x-axis). Different quantities of interest are considered: (A) *π*_11_, the proportion of SNPs associated with both traits; (B) *π*_10_, the proportion of SNPs associated with only the first trait; (C) *π*_11_/(*π*_11_+*π*_10_), the proportion of SNPs associated with the first trait that are also associated with the second trait; (D) Estimated parameters for the informative annotations.

### 3.2 Results of real data applications

#### 3.2.1 Real data and annotations

We applied our method to analyze 48 traits from 31 GWASs (Table S1). These traits span a wide range of phenotypes that include anthropometric traits (e.g. height and BMI), hematopoietic traits (e.g. MCHC and RBC), immune diseases (e.g. CD and IBD), and neurological diseases (e.g. Alzheimer’s disease and schizophrenia). We obtained summary statistics for these traits. We also obtained a total of 40 annotations based on genome-wide occupancy information of four histone marks (*H3K27me3*, *H3K36me3*, *H3K4me1* and *H3K4me3*) in 10 tissue groups (Blood/Immune, Adipose, Adrenal/Pancreas, Bone/Connective, Cardiovascular, CNS, Gastrointestinal, Liver, Muscle and Other) from the Roadmap Epigenomics Project (Roadmap Epigenomics Consortium, et al., 2015). Data processing details are provided in Text S2. We analyzed each of the 1,128 trait pairs with different methods. Methods considered include univariate analysis, gwas-pw, GPA and iMAP. For iMAP, we considered two different approaches. The first approach (iMAP-naïve) did not integrate any histone annotation while the second approach (iMAP) analyzed all histone annotations jointly with penalized selection. Because GPA can only examine a small number of histone annotations, to allow for a fair comparison, we applied GPA-select using the same screening procedure described in the simulations section and included all the selected histone annotations into the GPA model for a final analysis. For all methods, we declared association significance based on an estimated FDR of 0.1%.

#### 3.2.1 Integrative analysis by iMAP improves association mapping power

We computed the number of significant associations detected by different methods for each trait pair (Figure 3). These results are based on an estimated FDR of 0.05, which can be less stringent than the usual 5×10^−8^ p-value threshold that aims to control for an FWER of approximately 0.05. Consistent with simulations, iMAP detected more associations than any other methods. Specifically, iMAP identified an average of 658 associated SNPs across trait pairs (Figure 3A), while iMAP-naïve, GPA-select, gwas-pw and univariate analysis identified 580, 463, 232 and 358 associated SNPs, respectively. In addition, iMAP is ranked as the best method in terms of identifying the largest number of associated SNPs in 703 (62.3%) trait pairs out of the 1,128 total (Figure 3B-E), while iMAP-naïve, GPA-select, gwas-pw and univariate analysis were ranked as the best in 7 (0.6%), 217 (19.2%), 200 (17.7%) and 1 (0.09%) trait pairs, respectively. Among the associated SNPs, an average of 3.8% (25) of them have pleiotropic associations with both traits. The proportion of pleiotropic SNPs is similar whether annotations were accounted for or not; indeed, iMAP-naïve also identified an average of 2.9% (17) SNPs to have pleiotropic associations with both traits. The relatively small proportion of pleiotropic associations detected by iMAP is consistent with previous studies (Sivakumaran, et al., 2011). However, among biological related traits, the proportion of pleiotropic associations can be large (Figure S13). For example, among pairs of lipid traits (HDL, LDL, TC and TG), iMAP identified an average of 24.5% (522) SNPs associated with two traits. Similarly, iMAP-naïve identified an average of 19.9% (364) SNPs associated with two traits.

**Fig.3.**
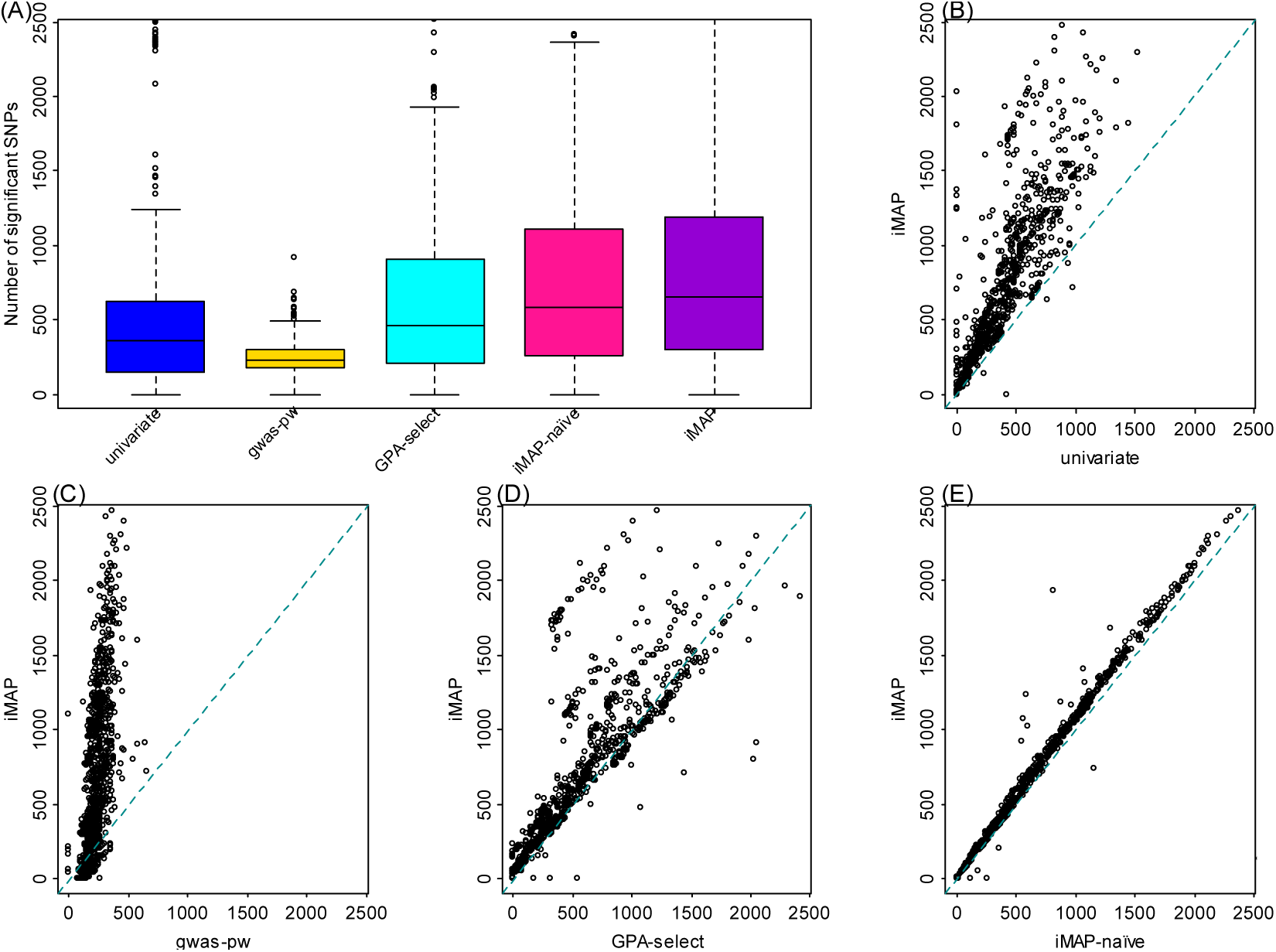
Power comparison among methods for the 1,128 trait pairs analyzed in the real data application. Methods for comparison include: univariate, gwas-pw, GPA-select, iMAP-naïve and iMAP. (A) Boxplots show the number of associated SNPs that pass the genome-wide significance threshold identified by various methods across 1,128 trait pairs. The number of associated SNPs identified by iMAP for each trait pair is also plotted against that identified by (B) univariate, (C) gwas-pw, (D) GPA-select and (E) iMAP-naïve.

As an illustrative example, we display a local genetic region that harbors a pleiotropic association for the trait pair of high-density lipoproteins (HDL) and triglycerides (TG). HDL and TG are blood metabolic phenotypes that are biologically related and have been previously shown to share a common genetic background (Bulik-Sullivan, et al., 2015; Pickrell, et al., 2016; Teslovich, et al., 2010). For the joint analysis of HDL-TG, iMAP identified a total of 2,305 associated SNPs, among which 376 (15.9%) of them are pleiotropic associations. The identified associated SNPs span a total of 51 loci (Table S4), among which 44 were previously known to be associated with either HDL or TG (MacArthur, et al., 2017). For illustration purpose, we display in Figure S14 a genomic region centered at a pleiotropic SNP, *rs1809167*, which resides near the gene *TRIB1* on 8q24.13. *TRIB1* is a protein-coding gene that is associated with multiple lipid traits and cardiovascular disease (MacArthur, et al., 2017; Soubeyrand, et al., 2016; Teslovich, et al., 2010). In the analysis, when we did not include SNP annotations, the PIP of *rs1809167* computed by iMAP-naïve is only 0.753 (Figure S14 A), below the genome-wide significance threshold (PIP=0.936, which corresponds to an FDR of 0.1%). After including SNP annotations, however, the PIP of the same SNP computed by iMAP increases to 0.988 (Figure S14 B), which passes the genome-wide significance threshold. Therefore, by integrating SNP functional annotations, iMAP has the potential to identify associations that may otherwise be missed by iMAP-naïve. In this analysis, iMAP also selected four histone annotations from three tissues to be relevant to the two traits, and these four annotations include Blood/Immune *H3K36me3* and *H3K4me1*, Bone/Connective *H3K36me3* and Liver *H3K4me1*. The estimated annotation coefficients of the four annotations are shown in Figure S14 B. Importantly, we find that the estimated effects of Liver *H3K4me1* and Bone/Connective H3K36me3 are larger than the other two annotations, suggesting that liver and Bone/Connective may play an important role in the biology of HDL and TG, consistent with previous findings (Kozlitina, et al., 2014; Lories, et al., 2013; Rivadeneira and Mäkitie, 2016; Roman, et al., 2015).

#### 3.2.2 iMAP identifies important histone annotations relevant to complex traits

Next, we examine the important histone annotations selected by iMAP across all trait pairs. iMAP selected at least one histone annotation for most pairs of traits examined (99.6%). On average, it selected 2.8 histone annotations (median=2.0) that influence the probability of a SNP to be associated with one trait, and selected 0.20 histone annotations (median=0) that influence the probability of a SNP to be associated with both traits. The small number of histone annotations selected in the later case is likely due to the relatively small number of pleiotropic SNPs and the subsequent low statistical power there. To examine individual histone annotations, for each histone annotation in turn, we counted the proportion of times that the histone annotation of interest was selected by iMAP across all analyzed trait pairs. In addition, we obtained the coefficient estimates for these selected histone annotations. Intuitively, if a histone annotation is an important predictor of SNP association, then it would be selected more often than the other nonimportant ones. In addition, it would have a higher estimated annotation coefficient than the others. We show both the coefficient estimates and the probability of being selected for the four histone annotations in Figure 4A. Specifically, across all tissues, *H3K36me3* and *H3K4me1* are selected more often than *H3K27me3* and *H3K4me3* (47.3% and 33.1% of the times versus 9.97% and 9.59% of the times). In addition, the estimated annotation coefficients of *H3K36me3* and *H3K4me1* are also larger than those of *H3K27me3* and *H3K4me3* (an average of 0.158 and 0.168 versus an average of 0.036 and 0.077). Both results are consistent with the important role of H3K36me3 and H3K4me1 in marking promoter or enhancer regions that are known to be predictive of SNP causality from previous univariate analyses (Roadmap Epigenomics Consortium, et al., 2015; The ENCODE Project Consortium, 2012).

**Fig.4.**
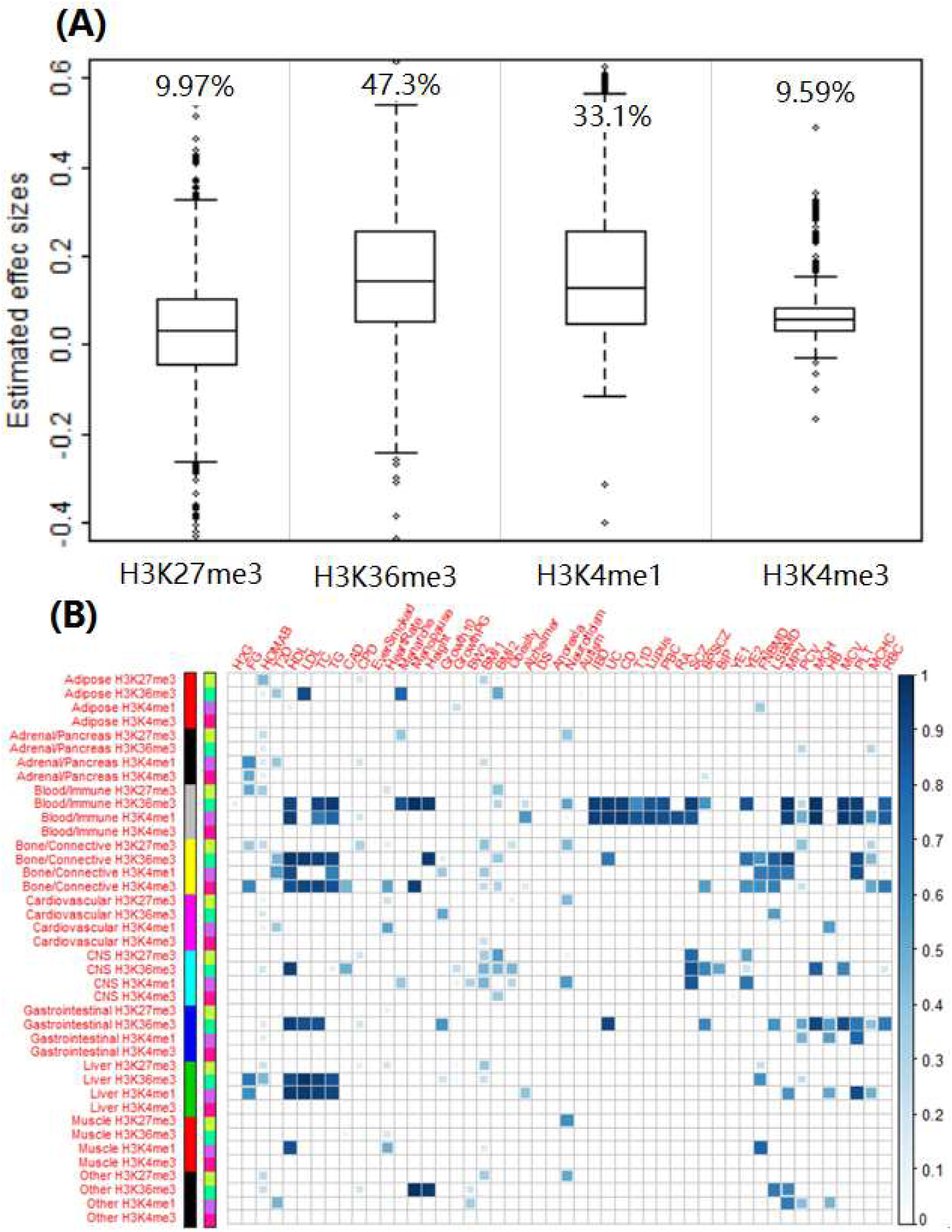
Annotation selection in the real data application. (A) Estimated annotation effect sizes for the selected annotations across all analyzed trait pairs. On top of the boxplots lists the percentage of times a histone annotation is selected across these trait pairs. (B) Relevance score for each annotation (rows) across all traits (columns). The relevance score quantifies the importance of an annotation for a particular trait of interest and is computed based on analysis of all trait pairs.

To further investigate trait-relevance of these histone annotations, for each trait/histone annotation pair in turn, we counted the proportion of times that the particular histone annotation was selected to have nonzero effects in all trait pairs that involve the trait of interest. The resulting value, which we simply refer to as the trait *relevance score*, is in the range of 0 and 1 and represents the relevance of the specific histone annotation to the given trait. Because histone annotations are tissue-specific, the resulting relevance scores also quantify trait-tissue relevance for trait tissue pairs. Results show that SNP associations in some traits are primarily predicted by histone annotations in a single tissue, while associations in other traits are predicted by histone annotations in a range of tissues (Figure 4B). For example, SNP associations in many immune diseases (e.g. UC, CD, IBD, T1D, Lupus, PBC and RA) are primarily related to two histone annotations, *H3K36me3* and *H3K4me1*, in the blood/immune tissue. Associations for most anthropometric traits (e.g. Height, BMI2, FNBMD, LSBMD, BMI1, BW2, G10, GPC and Obesity) can be predicted by marks in both the bone/connective and CNS tissues. Psychiatric disorders (e.g. SCZ, BPSCZ and BIP) are related to both blood/immune and CNS tissues. Lipids-related traits and diseases (e.g. HDL, LDL, TC, TG, CAD and X2hrGlucose) are related to blood/immune, bone/connective, gastrointestinal and liver tissues. While hematopoietic traits (e.g. MCHC, MCH, HB, MCV, MPV, PCV, PLT and RBC) are related to blood/immune, gastrointestinal and bone/connective tissues. Overall, the results are largely consistent with previous univariate analyses and reveal the complexity of trait-tissue relevance (Bradfield, et al., 2011; Liu, et al., 2015; Teslovich, et al., 2010).

#### 3.2.3 iMAP characterizes genetic relationship between pair of traits

Finally, we examine the estimated *π*_11_/(*π*_10_+*π*_11_) (or *π*_11_/(*π*_01_+*π*_11_)) values across all trait pairs. The proportion *π*_11_/(*π*_10_+*π*_11_) (or *π*_11_/(*π*_01_+*π*_11_)) can be asymmetric between any two traits and have been used to infer causality between the two traits (Pickrell, et al., 2016). The proportions estimated by iMAP are shown in Figure 5, and are largely consistent with the estimates by iMAP-naïve (Figure S15 A; Figure S15 B-C also show the proportions estimated by gwas-pw and GPA-select). In either case, the estimated proportions are largely symmetric, though with substantial asymmetric patterns. In particular, among all trait-pairs, 331 (29.3%) of them have an asymmetric pattern with the difference between the two proportions estimated to be larger than 10%. For example, the estimated proportions of SNPs associated with FG and HDL that are also associated with T2D are 0.983 and 0.507, respectively. On the other hand, the estimated proportions of SNPs associated with T2D that are also associated with FG and HDL are 0.568 and 0.058, respectively. The large proportion estimates between T2D and FG suggest that T2D and FG may share common biological pathways (Solovieff, et al., 2013). In contrast, the fact that SNPs associated with HDL often are also associated with T1D, but not vice versa, suggests that HDL may mediate SNPs effects onto T1D (Solovieff, et al., 2013). As another example, the estimated proportions of SNPs associated with the four blood lipid traits (HDL, LDL, TC and TG) that are also associated with CAD are high (0.669, 0.681, 0.815 and 0.724, respectively). In contrast, the proportions of SNPs associated with CAD that are also associated with the four blood lipid traits are small (0.068, 0.066, 0.064 and 0.096, respectively). The asymmetric relationship between lipid traits and CAD again suggests that lipid traits may mediate SNP effects onto CAD, consistent with previous findings (Bjornsson, et al., 2017; Pickrell, et al., 2016; Willer, et al., 2008).

**Fig.5.**
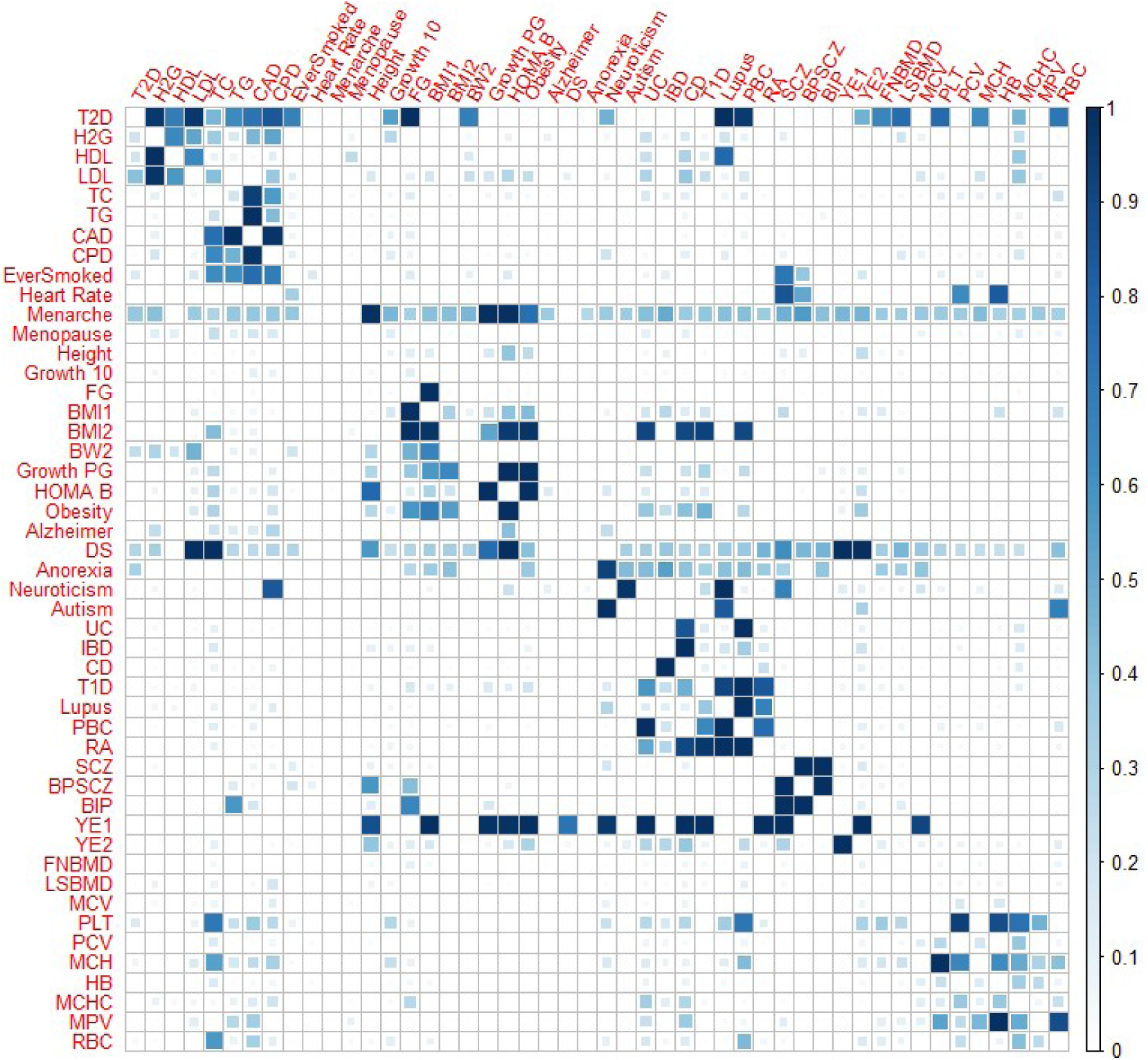
Estimated probability that a SNP associated with one trait (y-axis) is also associated with the other trait (x-axis), for 48 trait pairs in the real data application. Results are based on iMAP that performs annotation selection among 40 annotations.

## Discussion

We have presented a penalized Gaussian mixture model method, iMAP, that incorporates functional annotations for association mapping of pair-wise traits. iMAP properly accounts for phenotypic correlation, models association pattern across genome-wide SNPs, can accommodate both continuous and binary annotations, while capable of selecting informative annotations among a large set. iMAP is particularly useful for integrative analyses of GWAS summary statistics with a potentially large number of SNP annotations. While we have mainly focused on the use of Lasso (Tibshirani, 1996) for annotation selection, we have implemented the Elastic Net (Zou and Hastie, 2005) in the software to improve the performance of iMAP in the presence of correlated annotations. In addition, we have implemented the standard iMAP model without annotation input, so that the users can access the contribution of likelihood vs prior from annotation in the final association results. iMAP is also reasonably fast in the real data: it takes an average of 48 minutes (median = 45 minutes) to analyze a pair of traits in the presence of one annotation, and takes an average of 7.2 hours (median = 4.8 hours) when 40 annotations are used. With extensive simulations and real data application to 48 GWAS traits, we have illustrated the benefits of iMAP in terms of improving association mapping power, identifying important annotations, revealing genetic architecture underlying complex traits, as well as investigating relationship among phenotypes.

Throughout the text, we have used FDR thresholds to identify significant associations. While using FDR allows us to fairly compare the power of different methods based on the same threshold, using FDR also requires care in results interpretation for two reasons. First, compared to the commonly used p-value threshold of 5×10^−8^ that controls for an FWER of approximately 0.05, depending on the number of identified SNPs and the number of total SNPs, an FDR of 0.05 can be less stringent and can lead to the identification of unexpected many associated variants (Brzyski, et al., 2017). Consequently, the identified associations based on an FDR of 0.05 may suffer from lower replication rate as compared to those identified based on a p-value threshold of 5×10^−8^. Second, compared to FWER control, FDR control does not have the sub-setting property (Goeman and Solari, 2014). Specifically, because FWER controls for the probability of making a type I error, a FWER control at any level on the entire set of the identified SNPs guarantees the same (or more stringent) FWER level on any subset of the identified SNPs, regardless of SNP association evidence measured by p-value within the subset (Goeman and Solari, 2014). In contrast, FDR controls for the expected proportion of false discoveries, and an FDR control at any level on the entire set of the identified SNPs does not guarantee the same FDR level on any subset of the identified SNPs. In particular, SNPs with low PIP will have high local false discovery rate, and consequently, the subset of identified SNPs with low PIP will have higher FDR than what is controlled for on the entire set. Therefore, for FDR control, we recommend examining PIP as additional association evidence within the identified SNP set. We highlight these two important differences between FWER and FDR to help practitioners better interpret the results from iMAP as well as other Bayesian methods that rely on FDR for error control.

In the present study, we have mainly focused on the relatively simple task of analyzing pairs of traits. Extensions of iMAP to analyzing more than two traits may seem trivial conceptually, but present important statistical and computational challenges that need to be properly addressed. Specifically, with increasing number of traits (d), the number of possible association patterns increases exponentially (2^d^): a SNP can be associated with none of the traits, with any one trait, with any two traits,…, or with all traits. Unfortunately, direct and naïve modeling of the large number of possible association patterns would require an exponentially large number of ? parameters as well as an exponentially large number of annotation coefficients inside the mlogit model. Employing a large number of parameters not only would reduce the degrees of freedom and subsequently power, but also imposes computational hurdles (e.g. the computational complexity of iMAP also scales exponentially to the number of traits). Therefore, additional modeling assumptions on the possible association patterns are necessary to enable efficient and powerful analysis. Previous methods developed in eQTL mapping setting for identifying eQTLs in multiple tissues (Flutre, et al., 2013) have suggested that using either sparse (that a SNP is only associated with a small set of phenotypes) or group-structured (that there exist a number of phenotype groups, where a SNP is either associated with all phenotypes within the same group or associated with none of them) effect size assumptions are effective in modeling association patterns across tissues. Adapting these eQTL mapping methods to association mapping of pleiotropic effects is an interesting future avenue to explore.

Like previous other mixture methods (Chung, et al., 2014; Pickrell, et al., 2016), iMAP makes a key assumption that the joint likelihood for all SNPs is a simple product of the likelihoods from every SNP. However, because SNPs are in linkage disequilibrium (LD) and their genotypes are correlated with each other (Wall and Pritchard, 2003), assuming a simplified form of the joint likelihood is unrealistic. To examine whether iMAP remains effective under LD, we have performed a series of simulations using correlated SNPs instead of independent ones (Text S2). Briefly, we show that the power comparison results among methods are relatively stable with respect to LD and that iMAP still outperforms the other methods in identifying associated SNPs in the presence of LD. However, LD strongly influences the estimation of the proportions of causal SNPs in different association categories, and none of the methods examined here are accurate in estimating the causal proportions in the presence of LD (Text S2). We have attempted to improve the estimation of causal SNPs proportions by incorporating LD information as weights for the SNP likelihoods as suggested in (Liley, et al., 2017). However, we found that using either LD scores (Finucane, et al., 2015) or LDAK weights (Speed, et al., 2012) does not improve the accuracy in causal SNP proportion estimation. Besides the weighted likelihood, we have also attempted to prune SNPs before performing analysis for estimating causal SNP proportions (Kochi, et al., 2017). However, we found that the results from pruning can be unstable depending on how causal SNPs were simulated. Therefore, we view it as an important direction in the future to incorporate LD pattern into iMAP or other mixture model methods in a principled way. Such extension, together with efficient algorithmic innovations, will facilitate the accurate estimation of causal SNP proportion in various association categories and will likely further improve the power in fine mapping of pleiotropic traits (Kichaev and Pasaniuc, 2015; Kichaev, et al., 2014; Spain and Barrett, 2015).

## Acknowledgements

We thank all the GWAS consortium studies for making the summary data publicly available. The GWAS summary data source to these data consortium is given in the Supplementary Information.

